# Functional interactions between post-translationally modified amino acids of methyl-coenzyme M reductase in *Methanosarcina acetivorans*

**DOI:** 10.1101/771238

**Authors:** Dipti D Nayak, Andi Liu, Neha Agrawal, Roy Rodriguez-Carerro, Shi-Hui Dong, Doug A Mitchell, Satish K Nair, William W Metcalf

## Abstract

Methyl-coenzyme M reductase (MCR) plays an important role in mediating global levels of methane by catalyzing a reversible reaction that leads to the production or consumption of this potent greenhouse gas in methanogenic and methanotrophic archaea. In methanogenic archaea, the alpha subunit of MCR (McrA) typically contains four to six post-translationally modified amino acids near the active site. Recent studies have identified genes that install two of these modifications (thioglycine and 5-(S)-methylarginine), yet little is known about the installation and function of the remaining post-translationally modified residues. Here, we provide *in vivo* evidence that a dedicated SAM-dependent methyltransferase encoded by a gene we designated *mcmA* is responsible for formation of *S*-methylcysteine in *Methanosarcina acetivorans* McrA. Phenotypic analysis of mutants incapable of cysteine methylation suggests that the *S*-methylcysteine residue plays an important role in adaptation to a mesophilic lifestyle. To examine the interactions between the *S*-methylcysteine residue and the previously characterized thioglycine, 5-(S)-methylarginine modifications, we generated *M. acetivorans* mutants lacking the three known modification genes in all possible combinations. Phenotypic analyses revealed complex, physiologically relevant interactions between the modified residues, which alter the thermal stability of MCR in a combinatorial fashion that is not readily predictable from the phenotypes of single mutants. Surprisingly, high-resolution crystal structures of the various unmodified MCRs were indistinguishable from the fully modified enzyme, suggesting that interactions between the post-translationally modified residues do not exert a major influence on the physical structure of the enzyme, but rather serve to fine-tune the activity and efficiency of MCR.

## Introduction

Methyl-coenzyme M reductase (MCR) is an unusual and important enzyme, which to date has only been observed in strictly anaerobic, methane-metabolizing archaea (1, 2). In these organisms, MCR catalyzes the reversible inter conversion of methyl-coenzyme M (CoM, 2-methylmercaptoethanesulfonate) and coenzyme B (CoB, 7-thioheptanoylthreoninephosphate) to methane and a coenzyme B-CoM heterodisulfide:

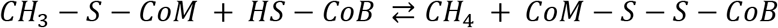

Although the methane-oxidizing activity of MCR in <u>**an**</u>aerobic <u>**me**</u>thane-oxidizing archaea (also referred to as ANME) leads to the consumption of *ca.* 90% of the methane produced in marine sediments (3, 4), the methanogenic activity of MCR dominates, leading to the net production of *ca.* 1 gigaton of methane annually (5). Accordingly, methanogenic archaea account for *ca.* 2/3rds of global methane emissions, with significant climate ramifications due to the fact that this abundant greenhouse gas has a warming potential 25 times higher than CO_2_ (5–7).

Although originally believed to be limited to a few taxa within the Euryarchaoeta (8), recent studies have broadened the diversity of MCR-encoding organisms to encompass all major phyla within the Archaea, including the Asgardarchaeaota, the closest known relatives to eukaryotes (9–12). MCR homologs in some of these uncultivated organisms are thought to be involved in the anaerobic catabolism of short-chain alkanes (1, 11, 13, 14). Thus, canonical MCRs lie within a larger family of ‘alkyl-coenzyme M reductases’ or ACRs (1, 11). Members of the larger ACR family play pivotal roles in archaeal evolution, climate change and the carbon cycle. They also offer new opportunities for the development of bio-based solutions for the production of methane and other renewable fuels (15). Unfortunately, challenges associated with the cultivation of strictly anaerobic methanogenic archaea, as well as with the *in vitro* activation of MCR have hampered functional characterization of this enzyme (1, 16). Thus, many fundamental properties of this important enzyme remain poorly understood.

MCR has a number of unusual traits (17–19). Crystallographic studies of MCR from methane-metabolizing archaea show a α_2_β_2_γ_2_ conformation for the protein complex (Figure 1A) (17–18). The complex possesses two active site pockets, each containing residues from both α-subunits. The active sites also contain tightly bound factor 430 (F_430_), a unique nickel porphinoid cofactor (20, 21) that, to date, has only been observed within MCR. The Ni(I) form of F_430_ is essential for catalysis; however the low reduction potential of the Ni(I)/Ni(II) couple (*ca.* −650 mV) renders F_430_ especially sensitive to oxidative inactivation, which, in turn, has substantially impeded *in vitro* biochemical characterization of MCR (16). The development of new experimental methods has allowed isolation of the fully active Ni(I) form of the enzyme, an especially challenging problem that required nearly twenty years to solve (1, 22). Recent studies of the fully active enzyme suggest that the reaction mechanism involves highly unusual methyl radical and Ni (II)-thiolate intermediate (23). Equally surprising was the discovery of between four to six rare post-translationally modified amino acids within the active site pocket, which eluded detection until high-resolution structures were obtained (18, 19, 24). These modified residues have been proposed to play roles in substrate binding, catalysis, or assembly of the MCR complex (24).

**Figure 1:**
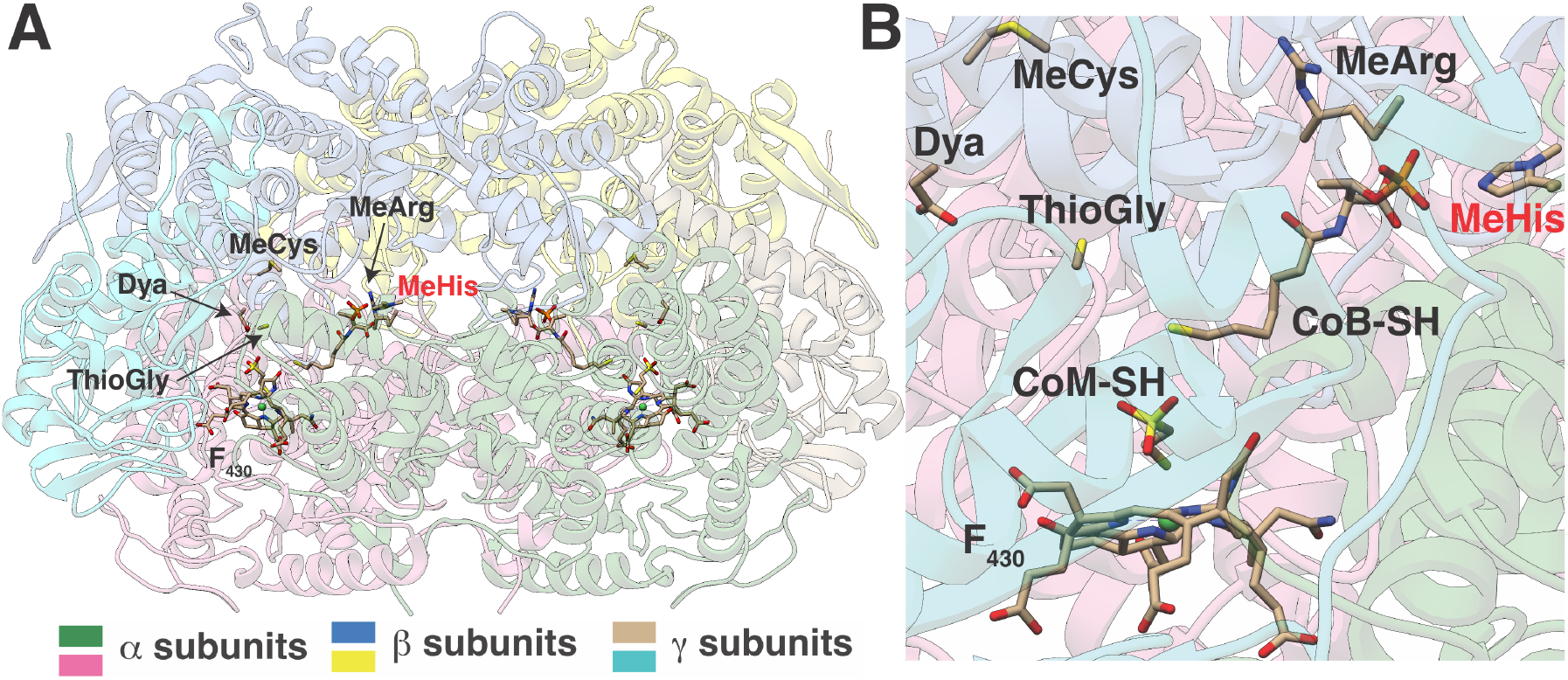
Crystal structure of MCR from *M. acetivorans*. **A)** The α_2_β_2_γ_2_ configuration of the MCR complex affinity-purified from *M. acetivorans* under aerobic conditions. The location of posttranslational modifications, the F430 cofactor, as well as coenzyme M (CoM-SH) and coenzyme B (coenzyme B-SH) are highlighted. No electron density corresponding to the affinity-tag was observed. **B)** A close-up of the active site within the McrA subunit in *M. acetivorans.* Dya: Didehydroasparate; MeCys: *S*-methylcysteine; MeArg: 5(S)-Methylarginine; MeHis: *N^1^-*Methylhistidine; ThioGly: Thioglycine. The red shading of the MeHis label indicates that this residue is found within the other α-subunit, illustrating the fact that residues from both α-subunits are found in each active site.

Mass spectrometric surveys of MCR from diverse methanogenic and methanotrophic archaea show that the modified residues in McrA fall into two broad classes: core and variable (24, 25). The core modified amino acids are widely conserved and include *N^1^*-methylhistidine (or 3-methylhistidine), 5-(S)-methylarginine, and thioglycine (Figure 1B). The widespread conservation of these core modifications is exemplified in the recently discovered genes responsible for installation of the thioglycine and 5-(S)-methylarginine modifications, which were originally designated as methanogenesis marker genes because of their universal occurrence in the genomes of sequenced methanogens (26–29). Given that 5-(S)-methylarginine and thioglycine are extremely rare in Nature, yet universally conserved in MCR, it was speculated that these residues must be essential for catalysis (24). Surprisingly, neither the *ycaO-tfuA* locus (MA0165/MA0164), responsible for installing thioglycine, nor the gene (MA4551) encoding the radical SAM methyltransferase responsible for installing 5-(S)-methylarginine is essential for methanogenic growth of *Methanosarcina acetivorans* (26, 27). Nevertheless, mutants lacking these genes display severe growth defect on substrates with low free energy yields (such as dimethyl sulfide or acetate) or when the cells are grown under stressful conditions (such as elevated temperatures or oxidative stress) (26, 27). Thus, the modifications are important for normal function of MCR and possibly required for methanogens to adapt to non-ideal environments. The gene(s) installing *N^1^*-methylhistidine have yet to be identified; therefore, the biochemical and physiological ramifications of this rare amino acid remain unexplored.

The variable modified amino acids include *S*-methylcysteine, 2-(S)-methylglutamine, and 6-hydroxytryptophan (24, 30–32). The phylogenetic distribution of these modifications is uneven, sometimes differing even between closely related strains. For instance, the didehydroaspartate modification is present in the McrA subunit from *Methanothermobacter marburgensis*, but absent in *Methanothermobacter wolfeii* (30). The factors underlying the phylogenetic distribution of variable modified amino acids are currently unknown; however, the non-overlapping occurrence of certain residues, such as *S*-methylcysteine and 6-hydroxytryptophan, suggests that they might be functionally analogous (31).

While previous studies have shown the importance of the core modified residues, the function of variable modifications has yet to be addressed. Similarly, the potential interactions between modified residues, as well as their combinatorial influence on MCR structure, stability and activity remain uncharacterized. Here, we identify the gene involved in the installation of *S*-methylcysteine, characterize the physiology of an *M. acetivorans* mutant incapable of this modification and assess the thermostability of MCR derived from this strain. We also report the generation and characterization of mutants lacking thioglycine, 5-(S)-methylarginine, and *S*-methylcysteine in all possible combinations, along with the high-resolution crystal structures of all eight MCR variants. Taken together, the data provide evidence of epistastic interactions between modified amino acids in MCR that impact the stability and function of this unusual and important enzyme.

## Results

### Crystal Structure of the wild-type MCR from *M. acetivorans*

We previously described the addition of an affinity tag that allowed for the facile purification of MCR from *M. acetivorans* cells (33). Aerobic purification and crystallization of the tagged protein from *M. acetivorans* (*Ma*MCR) allowed determination of the structure to a Bragg limit of 1.65 Å using phases calculated by molecular replacement. Relevant data reduction and refinement statistics are provided in Supplemental Table S1.

As expected from the high level of sequence conservation (90% identity), the three-dimensional structure of *Ma*MCR displays an architecture that is nearly identical to that of MCR from *Methanosarcina barkeri* (*Mb*MCR; PDB code: 1E6Y), with a root mean square (RMS) deviation of all Cα atoms of 0.24 Å. The protein complex crystallized as a α_2_β_2_γ_2_ assembly with one complex in the crystallographic asymmetric unit. Electron density for the affinity tag was not evident (Figure 1A). Notably, the active site of *Ma*MCR shows electron density features corresponding to the F_430_ cofactor, as well as for the coenzyme M and coenzyme B, showing that the affinity tag does not disrupt the association of these cofactors. Given the 1.65 Å resolution of the structure, clear electron density features indicative of modified amino acids are evident for the following residues of α subunit: *N*-methylation at His271, 5-(S)-methylation of Arg285, *S*-methylation of Cys472, and the presence of a thioamide bond at Gly465 (Figure 1B). Desaturation of the Cα-Cβ linkage at Asp470 to yield a didehydroasparate cannot be discerned at this resolution, but was confirmed by mass-spectrometry (see below). Therefore, we have included the didehydroasparate modification in the structural model.

### Identification of a methyltransferase mediating the *S-*methylcysteine modification of Cys472 in McrA from *M. acetivorans*

Based on its proximity to the structural genes encoding MCR in *M. acetivorans*, we hypothesized that a SAM-dependent methyltransferase belonging to the InterPro superfamily IPR29063 (MA4545; AAM07884.1; Q8THH2) would be involved in the post-translational methylation of one or more amino acids in McrA (Figure 2A) (24, 32). Significantly, phylogenetic profiling of the MA4545 locus revealed that homologs are absent in two hyperthermophilic methanogens (*Methanocaldococcus janaschiii* and *Methanopyrus kandleri*) that have been shown to lack *S-*methylcysteine within MCR (24). The phylogenetic tree of MA4545 is incongruent with both the MCR phylogeny and the reference phylogeny of archaea built with concatenated housekeeping genes (Figure 2B) (9, 11). Accordingly, it seems likely that a gene loss event and subsequent HGT event in the last common ancestor of the *Methanosarcinaceae* family is responsible for the shared ancestry of this locus between members of the *Methanosarcina* genus and the distantly related *Methanobacteriales* (Figure 2B).

**Figure 2:**
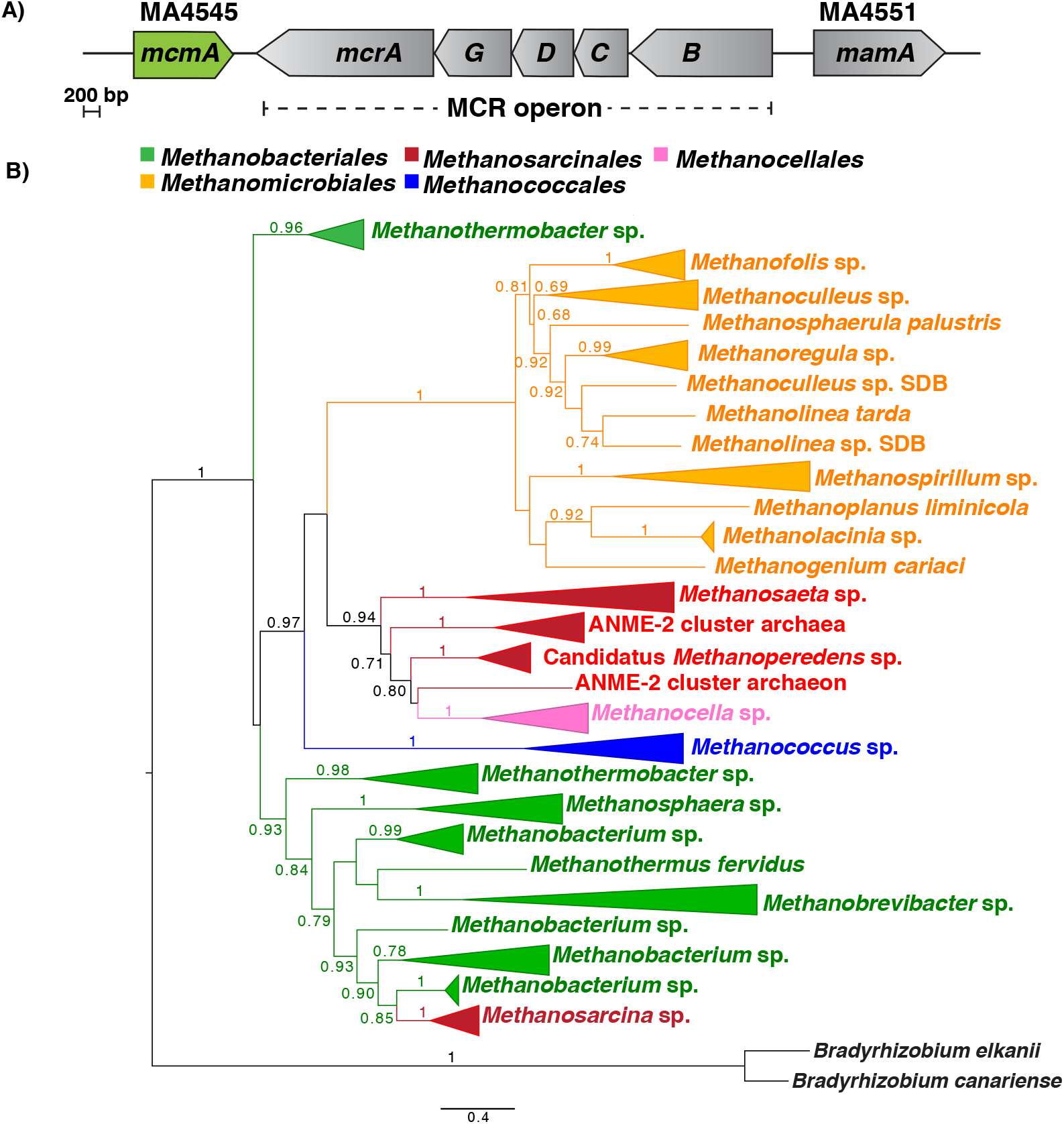
*In silico* analyses of McmA. **A)** Chromosomal organization of genes near the *mcr* operon in *Methanosarcina acetivorans.* Two *S*-adenosylmethionine (SAM) dependent methyltransferases are encoded on either side of the *mcr* operon. MA4551 encodes a radical SAM methyltransferase that was recently shown to be involved in the conversion of Arg285 in McrA to 5-(S)-methylarginine (27). We have named this gene *mamA* based on its proposed role (see text for details). **B)** A maximum-likelihood phylogenetic tree of the amino acid sequence of *mcmA* homologs in archaea. The node labels indicate support values calculated using the Shiomdaira-Hasegawa test using 1,000 resamples. Support values less than 0.6 have not been shown. The outgroup derives from bacterial McmA homologs (in black).

To test the hypothesis that MA4545 is responsible for *S-*methylation of Cys472 in McrA, we deleted the gene in *M. acetivorans* using recently developed Cas9-based genome editing tools (34). Matrix-assisted laser desorption/ionization time-of-flight mass spectrometry (MALDI-TOF-MS) showed that the McrA tryptic peptide containing the Cys residue of interest was 14 Da lighter in the mutant than in the corresponding peptide from wild-type, consistent with loss of a methyl group in the mutant (Figure 3C). The peptide was then subjected to high-resolution and tandem mass spectrometry, which confirmed the lack of Cys472 methylation in the mutant (Supplementary Figure S1). Taken together, these data clearly indicate that the MA4545 locus is involved in *S-*methylation of the Cys472 residue in McrA from *M. acetivorans.* Accordingly, we propose renaming this gene *mcmA* (<u>**m**</u>ethyl<u>**c**</u>ysteine <u>**m**</u>odification). Mass analysis of tryptic peptides containing the remaining modified amino acids showed that all were maintained in the *mcmA* mutant (Figure 3 and Supplementary Figure S2). Thus, installation of these modifications does not require the presence of methylcysteine.

**Figure 3:**
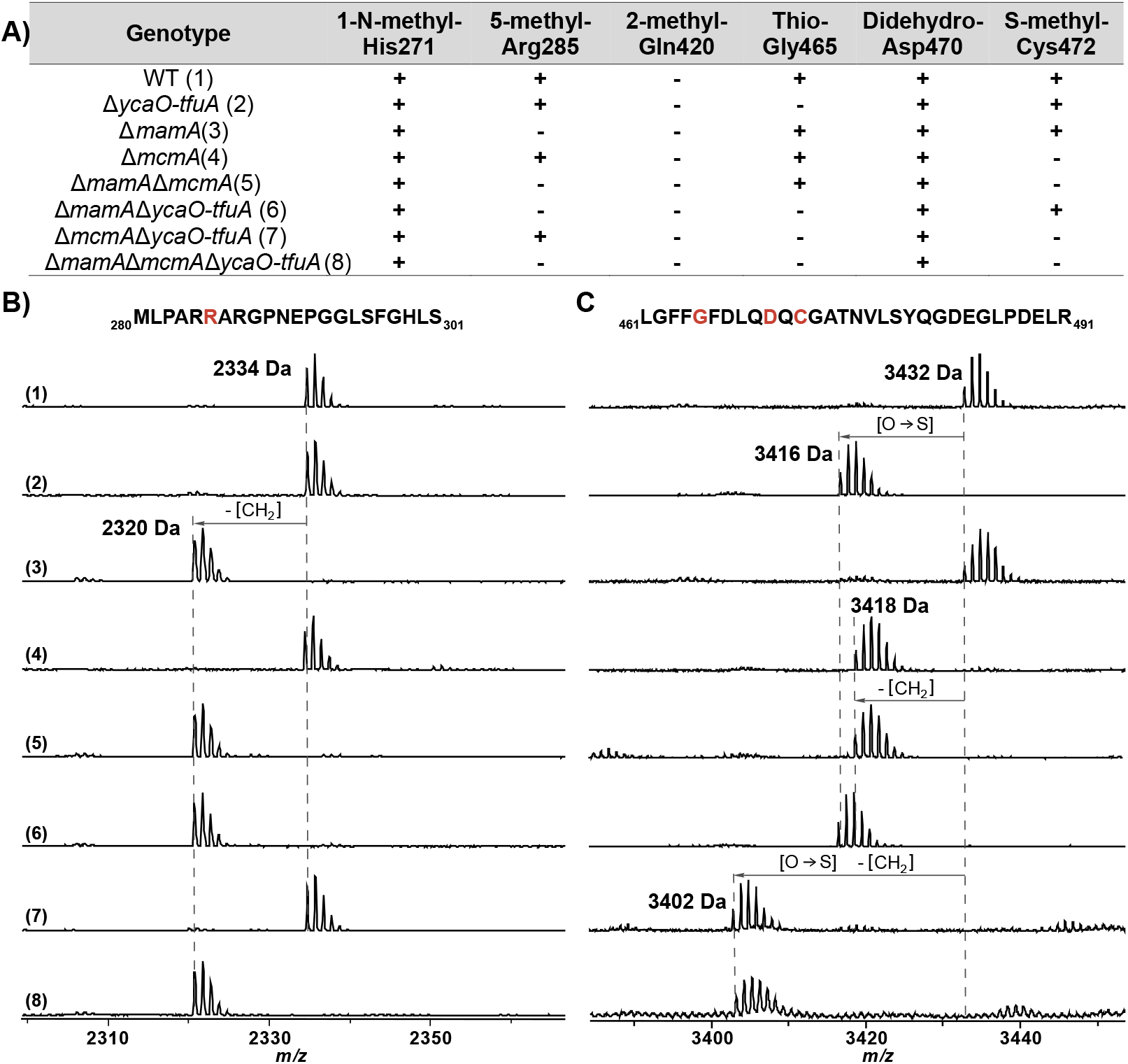
MALDI-TOF MS analysis of McrA. **A)** A list of all the posttranslational modifications found in McrA derived from the various mutants as indicated. **B)** Spectrum of the indicated peptides, which were obtained after AspN and GluC digestion of MCR from wild-type (WWM60) and mutants lacking *ycaO-tfuA, mcmA*, and *mamA* in all possible combinations. The M_280_-S_301_ peptide contains the Arg285 that is modified to 5-(S)-methylarginine by MamA. **C)** Spectrum of the indicated MCR tryptic peptide obtained from wild-type (WWM60) and mutants lacking *ycaO-tfuA, mcmA*, and *mamA* in all possible combinations. The L_461_-R_491_ peptide contains the Gly465 and Cys472 that are modified to thioglycine and *S*-methylcysteine by *ycaO-tfuA* and *mcmA*, respectively, as well as the didehydroaspartate at Asp470. Modified residues are red in the peptide sequence.

The ∆*mcmA* mutant was viable on all growth substrates tested (Figure 4A). Relative to wild-type, the mutant grew 30% slower on dimethyl sulfide (DMS) (p = 0.002 for an unpaired t-test with the means of three biological replicates) (Figure 4A). A 12% decrease in growth yield (measured as the maximum optical density at 600 nm) was observed on trimethylamine (TMA) (p = 0.018) (Figure 4B). Thus, even though the *S-* methylation of Cys472 in McrA is dispensable in *M. acetivorans*, it is clearly important for methanogenic growth on certain substrates. Curiously, the ∆*mcmA* mutant had better growth rates and yields than the wild type on some substrates, with more pronounced improvements at higher temperatures (Figure 4).

**Figure 4:**
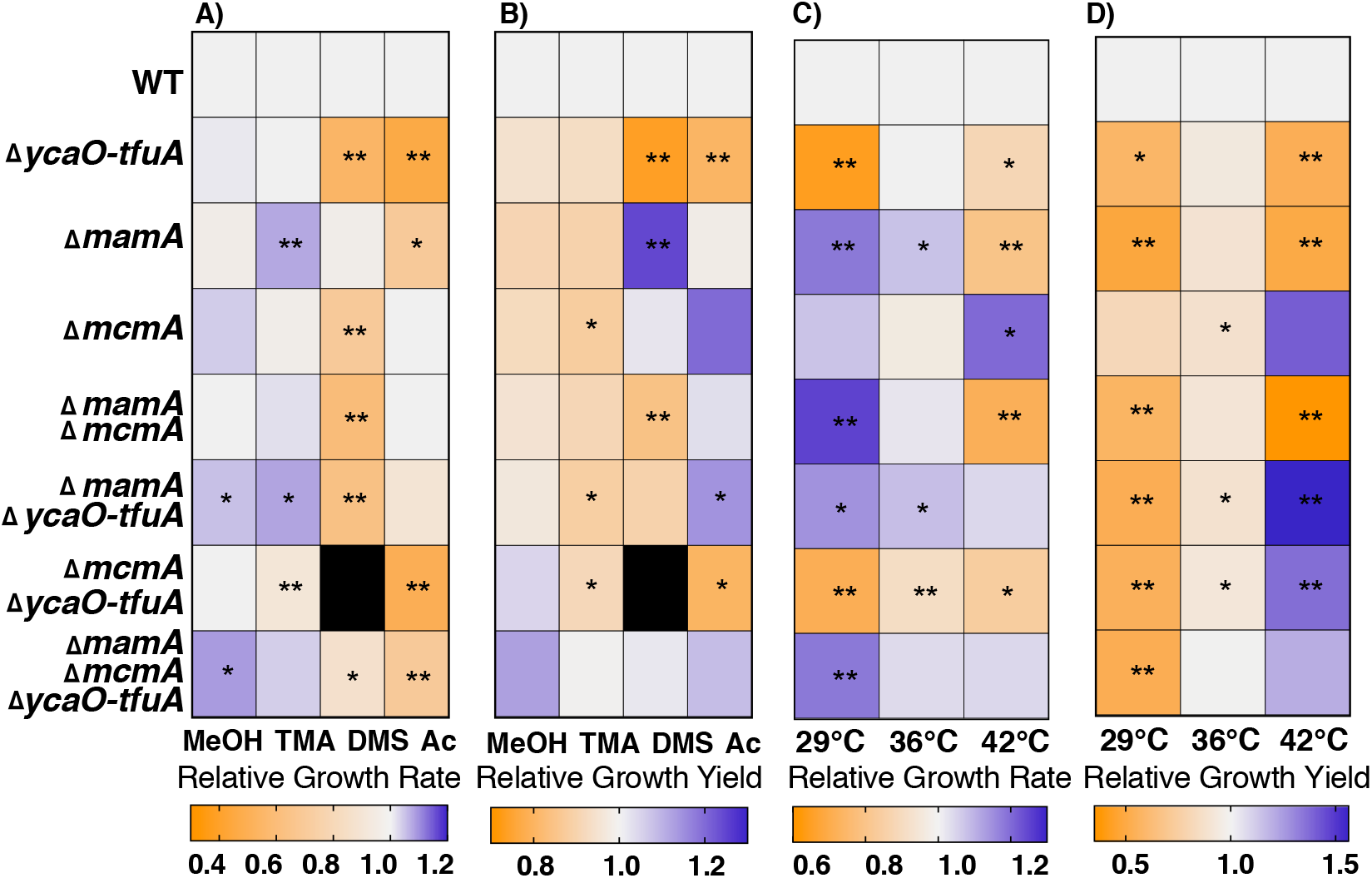
Phenotypic analyses on methanogenic substrates. Heat maps depicting **A)** growth rate or **B)** growth yield (measured as the maximum optical density at 600 nm) of mutants lacking *ycaO-tfuA, mcmA*, and *mamA* in all possible combinations relative to wild-type in bicarbonate-buffered High Salt medium supplemented with 125 mM methanol (MeOH), 50 mM trimethylamine hydrochloride (TMA), 20 mM Dimethylsulfide (DMS), or 40 mM sodium acetate (Ac) as the methanogenic substrates. All growth assays were performed at 36 °C. Heat maps depicting **C)** growth rate or **D)** growth yield (measured as the maximum optical density at 600 nm) of mutants in bicarbonate-buffered High Salt medium supplemented 50 mM TMA at three different temperatures as indicated. Statistically significant differences in growth parameters (p < 0.05 or p<0.01) relative to the wild-type as determined by a two-sided t-test are indicated with an * and ** respectively. The black box for the ∆*mcmA∆ycaO-tfuA* mutant in DMS supplemented HS medium indicates that no measurable growth was detected after six months of incubation. The primary data used to construct the heatmaps is presented in Supplementary Tables 2-13.

### Generation of combinatorial mutants of *M. acetivorans* lacking the thioglycine, 5-(S)-methylarginine, and *S-*methylcysteine modification in McrA

Since the α subunit of MCR contains multiple modified amino acids in spatial proximity (Figure 1B), epistasis between these residues may be important for optimal enzyme function. To test this, we generated deletion mutants lacking the genes responsible for installing *S*-methylcysteine, thioglycine and 5-(S)-methylarginine in all possible combinations (Supplementary Figure S3A). These include MA4551, which we have renamed *mamA* (<u>**m**</u>ethyl<u>**a**</u>rginine <u>**m**</u>odification) to better reflect its function (Supplementary Figure S4 and (27, 28)) and *ycaO-tfuA* (MA0165/MA0164), which coverts glycine to thioglycine (26, 29). The phenotypic analyses described below were carried out in mutants that encode wild-type MCR, while biochemical and structural studies were conducted with MCR purified from a second set of mutants that encode a modified *mcrG* allele (encoding the γ subunit of MCR) with an N-terminal a tandem-affinity purification tag comprising of a twin-Strep tag and a 3×FLAG tag (hereafter referred to as TAP tag) (Supplementary Figure S3B).

### Modified McrA residues from *M. acetivorans* are independently installed

Although the loss of single post-translational modifications does not effect the installation of other modifications (26, 27), the behavior of mutants lacking a combination of modified residues was unknown. To examine this, we identified all known post-tanslatioanlly modified amino acids in McrA from each of the mutants by MALDI-TOF-MS of peptides produced upon digestion with trypsin or a combination of the endoproteinases AspN and GluC (Figure 3). Additionally, high-resolution and tandem mass spectrometry techniques were employed to confirm the identity and location of each modification on the corresponding peptides (Supplementary Figures S5-S8). The modification state of McrA from *M. acetivorans* perfectly corresponds to the genotype of the mutant from which the protein was purified (Figure 3). These data validate the expected modification state of the mutants, while showing that the 5-(S)-methylarginine, thioglycine, and *S-*methylcysteine are independently installed.

### Growth analyses of combinatorial mutants reveal complex interactions between modified amino acids in McrA

Although direct interactions between the modified residues McrA are not apparent in the crystal structure, indirect interactions have been suggested to be important for enzyme catalysis (18, 19). For instance, both the thioglycine and *S-*methylcysteine are within van der Waals contact of Leu468, and such contacts are thought to cooperatively influence the local structure near the active site (Figure 1). To test whether these indirect interactions are physiologically relevant, and to uncover the nature of these interactions, we examined the growth phenotypes of mutants lacking the three modified residues in all possible combinations.

Growth analyses were performed on four different substrates: methanol (125 mM), TMA (50 mM), DMS (20 mM), and acetate (40 mM), which require substantially different methanogenic pathways and a wide range of free energy yields (∆G°’/mol CH_4_) (Supplementary Figure 9). Apart from the inability of the ∆*ycaO-tfuA/*∆*mcmA* double mutant to grow on DMS, all other mutants were viable on all the substrates tested (Figure 4A; Supplementary Tables S2-S13). Indeed, the ∆*ycaO-tfuA/*∆*mcmA* mutant had the most severe growth defect on all substrates tested, corroborating the hypothesis that the thioglycine and *S-*methylcysteine modifications interact synergistically (Figure 4A). Surprisingly, the triple deletion mutant grew faster than the ∆*ycaO-tfuA/*∆*mcmA* mutant, indicating that the unmodified Arg285 residue alleviates the growth defect observed in the absence of both the thioglycine and *S-*methylcysteine modifications (Figure 4A and 4B). On substrates with high free energy yields (methanol and TMA), most of the mutants lacking one or more of the modified residues grew as well as, or better than, the wild-type strain (Figure 4A). However, substantial growth rate defects, of varying magnitude depending on the strain, were observed on substrates with low free energy yields (DMS and acetate) (Figure 4A).

In a previous study, we observed that the ∆*ycaO-tfuA* mutant has a severe growth defect at elevated temperatures, which suggested that the thioglycine modification plays in a role in stabilizing the active site of MCR (26). Therefore, we assayed the growth phenotype of all mutants on 50 mM TMA at 29 °C, 36 °C, and 42 °C, where 36 °C represents the optimal growth temperature. While all mutants grew at every temperature, the growth phenotypes varied dramatically (Figure 4C, Figure 4D, Supplementary Tables S9-S13). Indeed, the mutant lacking all three modified residues grew 18% faster than wild-type at 29 °C (p = 0.001) (Figure 4C, Figure 4D, Supplementary Table 10). Significantly, the mutant lacking only the *S-*methylcysteine modification had a 20% increase in growth rate relative to wild-type at 42 °C (p = 0.018) (Figure 4C, Supplementary Table 11), indicating that this modification might be involved in the adaptation of MCR to growth under mesophilic conditions. In aggregate, the growth phenotypes of the double or triple mutants could not have been predicted by the studying the single mutants in isolation, which is a strong indicator of extensive, physiologically relevant, interactions between modified McrA amino acids.

### Interactions between thioglycine and 5-(S)-methylarginine are important for the in vitro thermal stability of MCR

The sign and magnitude of interactions between modified amino acids was especially evident during growth at elevated temperatures (Figure 4C and Figure 4D). To delve deeper into this phenotype, we investigated the global stability of the purified MCR complex using a SYPRO Orange-based thermofluor assay (35). Consistent with prior work (27), the melting temperature (T_m_) of MCR from the ∆*mamA* mutant was significantly lower (−6.5 °C; p< 0.001) than wild-type, whereas the T_m_ of the MCR from the ∆*mcmA* mutant was indistinguishable from wild-type (Figure 5A). Notably, the presence of Gly465 (as opposed to thioglycine) restored the T_m_ of MCR with an unmodified Arg285 residue, irrespective of whether Cys472 was modified (Figures 5B and 5C). Taken together, these data suggest that the 5-S-methylarginine and thioglycine modifications, as well as interactions between them, influence the thermal stability of MCR, again in ways that are not easily predicted from the stability of MCR lacking single modifications.

**Figure 5:**
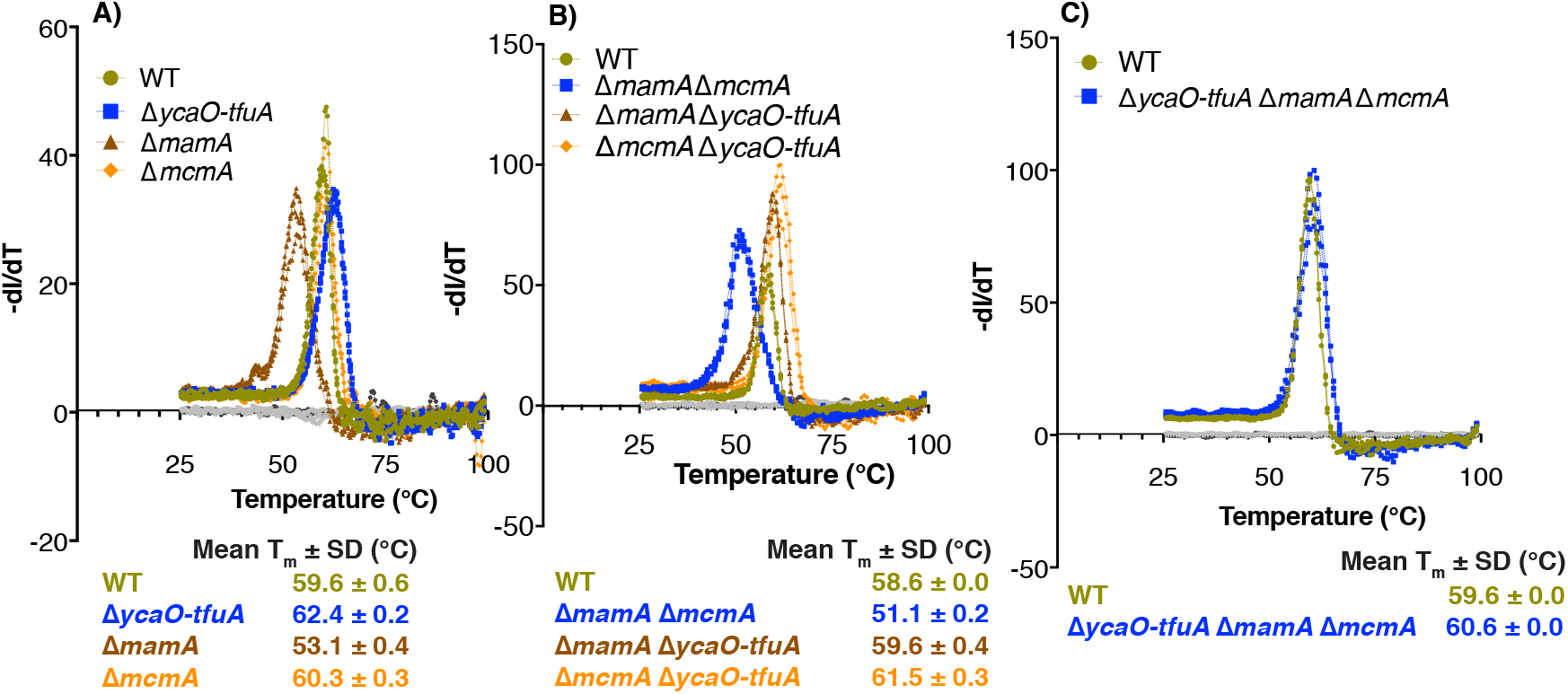
Melting temperature of the MCR complex. The thermal stability of the tandem-affinity tagged MCR complex purified from **A)** wild-type and single mutants lacking *ycaO-tfuA, mcmA*, or *mamA*, **B)** wild-type and double mutants lacking two of *ycaO-tfuA, mcmA*, and *mamA* in all possible combinations, and **C)** wild-type and the triple mutant lacking *ycaO-tfuA, mcmA*, and *mamA* was measured using the Sypro-Orange dye based thermofluor assay. The inflection point of the first differential curve for the fluorescence intensity relative to the temperature (-dI/dT) for each of three technical replicates was used to calculate the mean melting temperature (Tm) ± standard deviation (in °C) of the MCR complex. The no dye control (in grey) lacks the Sypro Orange dye and the no protein control (in black) was performed with elution buffer instead of purified protein.

### Structural analyses of combinatorial mutants

To gain molecular insights into the nature of the *S*-methylcysteine, thioglycine and 5-(S)-methylarginine interactions and their influence on the activity and stability of MCR, we solved the crystal structure of TAP-tagged MCR derived from each of the aforementioned mutants at resolutions between 1.9-2.3 Å (Supplementary Table 1). Crystals of the different variants occupy different unit cell settings, which rules out crystal packing artifacts as a potential source of bias between the structures. Examination of unbiased electron density maps reveals the presence and/or absence of the expected modified residues in the MCR variants obtained from the eight mutants.

The global structures of MCR variants purified from the combinatorial mutants remain unchanged relative to the wild-type, consistent with the lack of long-range interactions between the modified residues. Surprisingly, the structural analyses show that there are also no significant local changes to the active site pocket in any of the variants (Figure 6). Given the observation that the thioglycine and 5-(S)-methylarginine modifications influence thermal stability, we had expected the corresponding structures to reveal active site perturbations that would cause these differences. The similarity of the structures is also surprising given the extensive side chain interactions that occur at the site of each particular modification, which span the different subunits of MCR. We note, however, that because the crystal structures are likely to capture the lowest energy conformational state of each of the variant, dynamic movements that may occur at the voids created by removal of the amino acid modifications are unlikely to be captured in these static structures.

**Figure 6:**
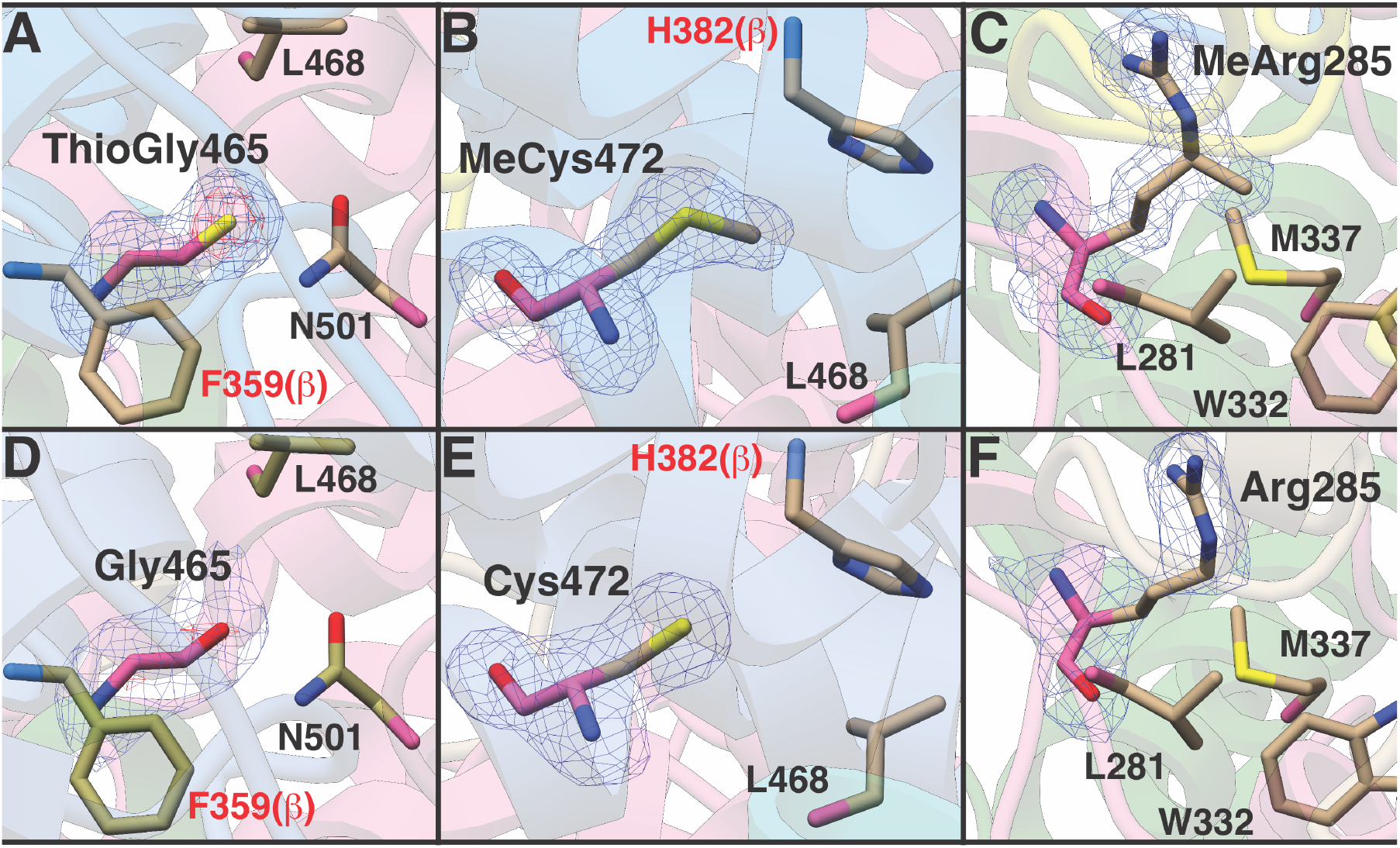
Absence of modified amino acids does not change the structure of MCR. **A), B), C)** The local structure near Gly465 (modified to thioglycine), Cys472 (modified to *S-*methylcysteine), and Arg285 (modified to 5-(S)-methylarginine) in wild-type. **D), E), F)** The local structure near the unmodified Gly465 residue, unmodified Cys472 residue, and unmodified Arg285 residue in MCR derived from the ∆*ycaO-tfuA, ∆mcmA, ∆mamA* single mutants respectively. The structure of MCR purified from the double and triple mutants was indistinguishable from that show here, with RMSD between all pairs ranging between 0.1-0.3 Å.

## Discussion

All characterized MCRs contain a set of core and variable modified amino acids near the active site (1, 9, 24). For the last two decades, researchers have speculated on the role of these modifications vis-à-vis the function of MCR. These hypotheses have ranged from certain modified residues being critical for activity to others playing minor roles (1). Recent studies have identified genes installing thioglycine and 5-(*S*)-methylarginine and shown that neither of these residues is essential for catalysis, although they might be important for the structural integrity of the MCR complex (26, 27). Here, we build on our previous study (26) by identifying a SAM-dependent methyltransferase involved in the installation of *S*-methylcysteine, a variable modified amino acid, and also uncovering evidence that epistasis between the three modified residues is also important for MCR *in vivo*.

The data presented above show that a dedicated SAM-dependent methyltransferase (renamed *mcmA*) is responsible for the *S-*methylation of a conserved Cys472 residue in McrA derived from *M. acetivorans* (Figure 3). Consistent with this conclusion, the C-terminus of McmA contains conserved residues that are involved in binding SAM. Thus, we suspect that the N-terminal domain of the protein, which lacks any identifiable domains, is involved in substrate recognition and/or binding. Homologs of McmA are especially widespread in methanogens that grow at moderate temperatures; however, they are absent from hyperthermophilic methanogens like *M. kandleri* and *M. janaschii*, as well as from psychrophilic methanogens like *Methanococcoides burtonii* or *Methanogenium frigidum* (Figure 2B). This phylogenetic distribution suggests that the *S-* methylcysteine modification is involved in the adaptation of MCR to mesophilic environments (Figure 2B). This idea is supported by our observation that the ∆*mcmA* mutant grew faster than the parent strain at both elevated (42 °C) and reduced (29 °C) temperatures (Figure 4C). Despite enhanced growth at high temperatures, the thermal stability of the MCR complex lacking the *S*-methylcysteine modification is indistinguishable from wild-type (Figure 5A). This indicates that the presence of *S-* methylcysteine does not improve the global stability of the enzyme complex. Furthermore, the structure of MCR from this mutant is essentially identical to that of the wild-type, including within the active site pocket near the modification site (Figure 6). Thus, we suspect that the temperature-dependent phenotypes of strains lacking *S*-methylcysteine are related to catalysis rather than structure. Curiously, homologs of McmA are absent in a few mesophilic methanogens, like the gut-associated *Methanomassiliicoccus* species (Figure 2B). Whether the *S-*methylcysteine modification is indeed absent in this organism, or whether another protein is capable of installing this modification, remains to be determined.

Despite the fact that the modified residues in MCR are in spatial proximity, only independent functions have been proposed for these unusual amino acids (*i.e.* they do not exert any influence on each other (22, 24). Our *in vivo* growth data, as well as the *in vitro* MCR thermal stability data, clearly show that each of the characterized modifications influences the function of the others. Moreover, our data demonstrate that the function of these modified amino acids is not reflected in static crystal structures. Thus, it seems highly likely that they exert their effects during enzyme turnover. Whether these effects are related to substrate binding and catalysis, or conformational/allosteric communication between the two active sites, as in the proposed two-stroke catalytic mechanism (1, 36), will await the *in vitro* characterization of active MCR from our mutants. Unfortunately, given its sensitivity to oxidative inactivation, *in vitro* kinetic characterization of MCR is especially problematic. The most widely used activation protocol relies hydrogenase activity (16). Unfortunately, as *M. acetivorans* does not produce active hydrogenases, this activation procedure is not directly transferable to our system. An alternate protocol involving pretreatment of cells with sodium sulfide to generate the Ni(III) from of the enzyme, followed by reduction of the purified enzyme with the strong reductant titanium citrate (37), has not been successful in our hands. Thus, kinetic characterization of the various unmodified MCR derivatives reported here currently remains out of reach.

Until a method for activation of *M. acetivorans* MCR is achieved, hypotheses for the functional roles of the modified MCR residues must rely on physiological data. MCR comprises *ca.* 10% of the total cellular protein and is often considered to be the rate-limiting step during methanogenic growth (23, 38). Therefore, if the loss of modifications alters MCR activity or the binding affinity for substrates, one would expect to see changes in growth rates and yields. For this reason, we assayed growth rates and yields on a variety of substrates that are used by distinct methanogenic pathways (Supplementary Figure 9). We note that a recent review suggested that anabolism is the rate-limiting step of methanogenic growth in batch cultures (1). While this is may be possible, it is difficult to rationalize this idea with the observation of faster (and slower) growth in the mutants described here, because MCR modification state is the only variable in these experiments and because MCR is not known to be involved in anabolism. Consistent with the idea that MCR is rate-limiting, we observed numerous, significant growth phenotypes. These include examples where the mutants performed significantly worse or significantly better than strains with fully modified MCR. The improved growth of modification-deficient mutants is especially surprising given the extensive conservation of post-translationally modified residues in MCR. Indeed, the triple mutant lacking thioglycine, *S*-methylcysteine and 5-(*S)*-methylarginine grew better than the parental strain under nearly every condition tested, which begs the question of why MCR is modified at all. A satisfactory answer to this question remains elusive.

The interactions between the modified residues are complex and difficult to predict from the phenotypes of double and single mutants; nevertheless, the observed phenotypes hint at possible mechanisms. The thioglycine and *S-*methylcysteine residues both interact with Leu468; whereas Leu468 and thioglycine also interact with Asn501, an important residue involved in coenzyme B binding (Figure 1B). Therefore, one hypothesis is that thioglycine and *S-*methylcysteine together facilitate the interaction between the thiol group of coenzyme B and the side-chain nitrogen of Asn501 to influence the K_m_ for coenzyme B. Likewise, the *S-*methylcysteine is in close proximity to His382 from McrB at the intersubunit interface and might similarly affect the binding of coenzyme B. This hypothesis is supported by the observation that the most severe growth defects are seen in mutants lacking *S*-methylcysteine and thioglycine. Curiously, our growth data also reveal that, despite being >14 Å away from Gly465 or Cys472, the unmodified Arg285 mitigates the deleterious effects of these unmodified amino acids (Figure 4). As each of the modified residues are involved in contacts within and across different subunits of MCR, we speculate that removal of the modifications has minimal effects on the structure. Essentially, deletion of the post-translational modification creates voids in the central core of MCR, adjacent to the active site, which compromises the catalytic stability of the enzyme during catalysis without changing the lowest entropy conformational state that is captured by X-ray crystallography.

Epistasis between modified residues was also evident when we characterized the global stability of the MCR complex *in vitro.* Among the three modified amino acids only 5-(*S*)-methylarginine plays a significant role in mediating the thermal stability of MCR (Figure 5A). Neither thioglycine nor *S*-methylcysteine influence thermal stability in isolation, but the unmodified Gly465 residue compensates for the deleterious impact of an unmodified Arg285 on global stability. A comparison of the crystal structures of the variant lacking 5-methylation on Arg285, with that lacking both this modification as well the thioglycine failed to reveal any notable changes in the active site that may account for the compensatory effects observed on thermal stability. Hence, it is likely that the effect of the mutations occurs either on the unfolded state of MCR or on transient intermediates formed during the folding process, without affecting the lowest energy ground state observed in the crystal structures.

In summary, recent studies have changed our view of the modified amino acids in MCR from biochemical novelties to evolutionary spandrels: features that are an offshoot of adaptation rather than a direct product thereof. Despite this paradigm shift, it is still perplexing as to why members of the ACR family contains between four to six unique and rare post-translational modifications. If the role of these modifications is to fill voids or contort the amino acid backbone, why didn’t the twenty standard amino acids suffice? It is entirely feasible that the functions of these modifications extend beyond the scope of conditions that can be tested in a laboratory setting. As we sample diverse groups of methanogenic archaea, ANMEs, and even anaerobic alkane oxidizing archaea and identify the pattern of modifications in their ACRs, maybe the underlying reasons will become apparent. In the near future, we expect that broader surveys of diverse ACRs coupled with laboratory-based genetic experiments will enable the design of appropriate experiments to tease apart the role of these unusual modified amino acids.

## Materials and Methods

### Phylogenetic analyses

One hundred closest homologs were extracted from the NCBI non-redundant protein database using the McmA amino acid sequence (MA4545; AAM07884.1; Q8THH2) or the MamA amino acid sequence (MA4551, AAM07890; Q8THG6) as queries in BLAST-P searches. Any partial sequences (i.e. sequences < 300 amino acids in length) were discarded and the full-length protein sequences were aligned using the MUSCLE plug-in (39) with default parameters in Geneious version R9 (40). Approximate maximum-likelihood trees were generated using FastTree version 2.1.3 SSE3 using the Jones-Taylor-Thornton (JTT) model + CAT approximation with 20 rate categories. Branch support was calculated using the Shimodaira-Hasegawa (SH) test with 1,000 resamples. Trees were displayed using Fig Tree v1.4.3 (http://tree.bio.ed.ac.uk/software/figtree/).

### Construction of gene-editing plasmids

All mutations in *M. acetivorans* were introduced using a Cas9-based genome editing technique (34). Mutagenic plasmids were derived from pDN201, a base vector containing Cas9 from *Streptococcus pyogenes* under the control of the tetracycline-inducible pMcrB(tetO1) promoter. A double-stranded DNA fragment, synthesized as a gblock (Integrated DNA Technologies, Coralville, IA, USA), expressing one or more single guide (sg) RNAs targeting the locus of interest was introduced at the *AscI* site in pDN201 using the HiFi assembly kit (New England Biolabs, Ipswich, MA, USA). Subsequently, a homology repair template, either generated by PCR amplification or synthesized as a gblock (Integrated DNA Technologies, Coralville, IA, USA), was introduced at the *PmeI* site in the sgRNA-containing vector using the HiFi assembly kit (New England Biolabs, Ipswich, MA, USA). A cointegrate of the resulting plasmid and pAMG40 (containing the pC2A replicon) was generated using the BP clonase II enzyme master mix (Thermo Fisher Scientific, Waltham, MA, USA) to enable autonomous replication of the mutagenic vector in *M. acetivorans.* All primers used to generate and verify plasmids are listed in Supplementary Table S15 and the plasmids used in this study are listed in Supplementary Table S16. Standard techniques were used for the isolation and manipulation of plasmid DNA. All pDN201-derived plasmids were verified by Sanger sequencing at the Roy J. Carver Biotechnology Center, University of Illinois at Urbana-Champaign and all pAMG40 cointegrates were verified by restriction endonuclease analysis.

### *In silico* design of sgRNAs for gene editing

All target sequences used for Cas9-mediated genome editing in this study are listed in Supplementary Table S17. Target sequences were chosen using the CRISPR site finder tool in Geneious version R9. The *M. acetivorans* chromosome and the plasmid pC2A were used to score off-target binding sites.

### *E. coli* growth and transformations

WM4489, a DH10B derivative engineered to control copy-number of oriV-based plasmids (41), was used as the host strain for all plasmids generated in this study (Supplementary Table S16). Electrocompetent cells of WM4489 were generated as described in (41) and transformed by electroporation at 1.8 kV using an *E. coli* Gene Pulser (Bio-Rad, Hercules, CA). Plasmid copy number was increased dramatically by supplementing the growth medium with sterile rhamnose to a final concentration of 10 mM. The growth medium was supplemented with a sterile stock solution of chloroamphenicol to a final concentration of 10 µg/mL and/or kanamycin to a final concentration of 25 µg/mL as appropriate.

### Transformation of *M. acetivorans*

All *M. acetivorans* strains used in this study are listed in Supplementary Table S18. Liposome-mediated transformation was used for *M. acetivorans* as described previously (42) using 10 mL of late-exponential phase culture of *M. acetivorans* and 2 µg of plasmid DNA for each transformation. Puromycin (CalBiochem, San Diego, CA) was added to a final concentration of 2 µg/mL from a sterile, anaerobic stock solution to select for transformants containing the *pac* (puromycin transacetylase) cassette. The purine analog 8-aza-2,6-diaminopurine (8ADP) (R. I. Chemicals, Orange, CA) was added to a final concentration of 20 µg/mL from a sterile, anaerobic stock solution to select against the *hpt* (phosphoribosyltransferase) cassette encoded on pC2A-based plasmids. Plating on HS medium containing 50 mM TMA solidified with 1.7 % agar was conducted in an anaerobic glove chamber (Coy Laboratory Products, Grass Lake, MI) as described previously (43). Solid media plates were incubated in an intra-chamber anaerobic incubator maintained at 37 °C with N_2_/CO_2_/H_2_S (79.9/20/0.1) in the headspace as described previously (43).

### Growth assays for *M. acetivorans*

*M. acetivorans* strains were grown in single-cell morphology (44) in bicarbonate-buffered high salt (HS) liquid medium containing one of the following: 125 mM methanol, 50 mM trimethylamine hydrochloride (TMA), 40 mM sodium acetate, or 20 mM dimethylsulfide (DMS). Most substrates were added to the medium prior to sterilization. DMS was added from an anaerobic stock solution maintained at 4 °C immediately prior to inoculation. For growth analyses, 10 mL cultures were grown in sealed Balch tubes with N_2_/CO_2_ (80/20) at 8-10 psi in the headspace. Growth measurements were conducted with three independent biological replicates derived from colony-purified isolates. Each replicate was acclimated to the growth substrate or growth temperature for a minimum of five generations prior to quantitative growth measurements. A 1:10 dilution of late-exponential phase cultures was used as the inoculum for growth analyses.

### Affinity purification of MCR for MS analyses and Thermofluor assays

TAP-tagged MCR was purified from 250 mL of late-exponential phase culture grown in HS + 50 mM TMA at 36 °C. Protein purification was performed under aerobic conditions and cells were harvested by centrifugation (3,000 × *g*) for 15 min at 4 °C. The cell pellet was lysed in 10 mL salt-free wash buffer (50 mM NaH_2_PO_4_, pH = 8.0) and sodium chloride was added to a final concentration of 300 mM post-lysis. The cell lysate was treated with DNase and clarified by centrifugation (17,500 × *g*) for 30 min at 4 °C. The supernatant fraction was loaded on a column containing 1 mL Streptactin Superflow Plus slurry (50% suspension; QIAGEN, Germantown, MD, USA) equilibrated with 8 mL of the wash buffer (50 mM NaH_2_PO_4_, 300m M NaCl, pH = 8.0). The column was washed four times with 2 mL of Wash buffer and the purified protein was eluted in four fractions with 0.5 mL Elution buffer (50 mM NaH_2_PO_4_, 300 mM NaCl, 50 mM biotin, pH = 8.0) per fraction. The protein concentration in each fraction was estimated using the Coomassie Plus (Bradford) assay kit (Pierce Biotechnology, Thermo-Scientific, Rockford, IL, USA) with BSA (bovine serum albumin) as the standard per the manufacturer’s instructions. The highest protein concentration (*ca.* 1 mg/mL) was obtained in fraction 2 therefore this fraction was used to conduct the Thermofluor assay. To visualize the purified protein, 10 µL of fraction 2 was mixed with an equal volume of 2× Laemmli sample buffer (Bio-Rad, Hercules, CA) containing 5% β-mercaptoethanol, incubated in boiling water for 10 min, loaded on a 4-20% gradient Mini-Protean TGX denaturing SDS-PAGE gel (Bio-Rad, Hercules, CA) and run at 70 V until the dye-front reached the bottom of the gel. The gel was stained using the Gel Code Blue stain reagent (Thermo Fisher Scientific, Waltham, MA) as per the manufacturer’s instructions.

### Proteolytic digestion of purified MCR

For trypsinolysis, 300 µg of purified MCR was digested with mass spectrometry (MS)-grade trypsin (1:100 *w*/*w* ratio; Thermo-Scientific, Rockford, IL, USA) at a final MCR concentration of 1 mg/mL in 50 mM NH_4_HCO_3_ at 37 °C for 14 h. The tryptic peptides were desalted using a C-18 zip-tip using aqueous acetonitrile or further purified by high performance liquid chromatography (HPLC) prior to MS analysis. For AspN and GluC double digestion, 50 µg of purified MCR was digested with endoproteinase GluC (1:200 *w*/*w* ratio; New England BioLabs, Ipswich, MA, USA) and endoproteinase AspN (1:200 *w*/*w* ratio; Promega, Madison, WI, USA) at a final MCR concentration of 1 mg/mL with the addition of 20% AspN 1× buffer (New England BioLabs, Ipswich, MA, USA) and GluC 2× buffer (New England BioLabs, Ipswich, MA, USA) at 37 °C for 24 h. The resulting digested peptides were desalted with C-18 zip-tips using acetonitrile with 0.1% formic acid prior to MS analysis.

### HPLC purification of MCR tryptic fragments

The aforementioned tryptic peptides were subjected to HPLC analysis using a C18 column (Macherey-Nagel, 4.6 × 250 mm, 5 µm particle size). Acetonitrile and 0.1% (*v*/*v*) formic acid were used as the mobile phase. A linear gradient of 20 to 80% acetonitrile over 23 min at 1 mL/min was used to separate the peptides. Fractions were collected at one-minute intervals and analyzed by matrix-assisted laser desorption/ionization time-of-flight mass spectrometry (MALDI-TOF-MS) with alpha-cyano-4-hydroxycinnamic acid (CHCA) as matrix using a Bruker UltrafleXtreme instrument (Bruker Daltonics, Billerica, MA, USA) in reflector positive mode at the University of Illinois School of Chemical Sciences Mass Spectrometry Laboratory. The fragment of interest elutes between 11-12 min regardless of the modification. The fraction was dried under vacuum using a Speedvac concentrator (Thermo Fisher Scientific, Waltham, MA, USA) for further analysis.

### MS analysis of MCR peptidic fragments

MALDI-TOF-MS analysis was performed on the desalted MCR fragments as described above. For high-resolution electrospray ionization (ESI) MS/MS, samples were dissolved in 35% acetonitrile and 0.1% formic acid and directly infused using an Advion TriVersa Nanomate 100 into a ThermoFisher Scientific Orbitrap Fusion ESI-MS. The instrument was calibrated weekly, following the manufacturer’s instructions, and tuned daily with Pierce LTQ Velos ESI Positive Ion Calibration Solution (Thermo Fisher Scientific, Waltham, MA). The MS was operated using the following parameters: resolution, 100,000; isolation width (MS/MS), 1 *m/z*; normalized collision energy (MS/MS), 50; activation q value (MS/MS), 0.4; activation time (MS/MS), 30 ms. Data analysis was conducted using the Qualbrowser application of Xcalibur software (Thermo Fisher Scientific, Waltham, MA, USA). HPLC-grade reagents were used to prepare samples for mass spectrometric analyses.

### Thermofluor Assay

A 5000× concentrate of SYPRO Orange protein gel stain in Dimethylsulfoxide (DMSO) (Thermo Fisher Scientific, Waltham, MA, USA) was diluted with the elution buffer to generate a 200× stock solution. Purified MCR from fraction 2 was diluted to *ca.* 500 µg/mL with the elution buffer. 5 µL of the 200× stock solution of SYPRO Orange protein gel stain was added to 45 µL of purified protein with a concentration of *ca.* 500 µg/mL in 96-well optically clear PCR plates. A melt curve was performed on a Mastercycler ep realplex machine (Eppendorf, Hamburg, Germany) using the following protocol: 25 °C for 30 s, ramp to 99 °C over 30 min, return to 25 °C for 30 s. SYPRO Orange fluorescence was detected using the VIC emission channel and the temperature corresponding to the inflection point of the first derivative (-dI/dT) was determined to be the melting temperature (T_m_). Appropriate no-dye controls and no-protein controls were examined and each sample was run in triplicate. The Thermofluor assay was conducted within 12 h of protein purification.

### Affinity purification of MCR from *M. acetivorans* for crystallography

TAP-tagged MCR was purified as described above for the MS analyses and Thermofluor assays with the following modifications. Protein was purified under aerobic conditions from 500 mL of late-exponential phase culture grown in HS + 50 mM TMA at 36 °C. The crystallography wash buffer comprised of 100 mM Tris.HCl, 300 mM NaCl and 2 mM dithiothreiotol (DTT) at pH 8. The crystallography elution buffer contained 2.5 mM desthiobiotin in addition to the other components of the crystallography wash buffer. Four 250µL fractions of purified MCR were collected and visualized using a 12% Mini-Protean TGX denaturing SDS-PAGE gel (Bio-Rad, Hercules, CA).

### Crystallization of MCR from *M. acetivorans*

MCR was concentrated to 25 mg/ml prior to crystallization via ultrafiltration. The crystallization trials utilized 2 μl sitting drops composed of 0.9: 0.9: 0.2 (protein: reservoir solution: additive screen) that were equilibrated against a 500 μl volume of the reservoir solution at room temperature. The reservoir solution contained 0.2 M ammonium acetate, 0.1 M sodium acetate, pH = 4 and 15 % (w/v) PEG 4000. Similarly, the MCR variants were concentrated to 3-30 mg/ml. The crystals were obtained in a hanging drop format composed of 2 μl mixture of 0.9: 0.9: 0.2 (protein: reservoir solution: additive screen) equilibrated against the same reservoir solution as the wild-type. Prior to freezing by vitrification in liquid nitrogen, crystals were soaked in reservoir solution supplemented with an additional 20 % (v/v) glycerol or 30 % (w/v) PEG 4000.

## Acknowledgements

We thank Keith Brister and the staff at LS-CAT (Argonne National Labs) with facilitating data collection. The authors acknowledge the Division of Chemical Sciences, Geosciences, and Biosciences, Office of Basic Energy Sciences of the U.S. Department of Energy through Grant DE-FG02-02ER15296 to WWM, and the National Institutes of Health (GM097142 to D.A.M.) for funding this work. DDN was supported by a Carl R. Woese Institute for Genomic Biology postdoctoral fellowship and the Simons Foundation Life Sciences Research Foundation postdoctoral fellowship. AL was the recipient of the Alice Helm Graduate Research Excellence Fellowship in Microbiology from the Department of Microbiology at the University of Illinois at Urbana-Champaign. The Bruker UltrafleXtreme MALDI TOF/TOF mass spectrometer was purchased in part with a grant from the National Institutes of Health (S10 RR027109 A).

